# Mechanistic insights into the role of Ca^2+^-stimulated AMPK in the secretion of cellulases during carbon stress

**DOI:** 10.1101/2023.01.16.524192

**Authors:** Anmoldeep Randhawa, Tulika Sinha, Maitreyee Das, Olusola A. Ogunyewo, Kamran Jawed, Syed Shams Yazdani

**Author notes:** Corresponding author: Syed Shams Yazdani, Microbial Engineering Group, International Centre for Genetic Engineering and Biotechnology, New Delhi 110067.

## Abstract

The response of filamentous fungi towards recalcitrant carbohydrates is majorly governed by transcriptional activators of cellulase genes; however, little is known about the downstream events beyond transcription. We show here in *Penicillium funiculosum* that increasing the expression of a transcriptional activator CLR-2 in the catabolically derepressed strain, ΔMig1, didn’t exhibit a synergistic effect on cellulase production unless Ca^2+^ was simultaneously increased. The RNA-seq screen for Ca^2+^-activated kinases identified SNF1-AMPK and SSP1-AMPKK as being specific to cellulose induction. Deletion of *snf1* led to negligible secretion of cellulase upon induction. Quantitative whole-cell proteomics followed by chemical-genetic experiments with *snf1-*deleted strain showed that Ca^2+^-signaling channelizes carbon, nitrogen and energy sources towards cellulase production. Further, Ca^2+^-signaling phosphorylates SNF1-AMPK via SSP1, which in turn downregulates the phospho-HOG1 levels, leading to stimulus for cellulase secretion during carbon stress. The findings reported here are significant for understanding fungal pathology and developing second-generation biorefineries.

## Introduction

The ability of pathogenic fungi to survive extreme conditions as saprophytes accounts for the rising cases of emerging fungal infections of plants, animals, and humans in 21^st^ century. The ability to hydrolyze recalcitrant structural polysaccharides of plant cell walls is an important survival factor for the phytopathogenic as well as for the saprophytic fungi. Expression of enzymes hydrolyzing these alternate carbon sources is repressed by glucose and other fermentable sugars, and is stringently activated in a substrate-specific manner during carbon stress^1^. Recent studies have identified both positive and negative regulatory circuits governing cellulase production at transcriptional level^2–5^.

A global negative regulator of secondary carbon metabolism, MIG1 (also called catabolite repressor CRE-1), controls transcriptional dynamics of cellulolytic genes^6–10^. Deletion of *mig1* gene results in catabolically de-repressed strains of filamentous fungi^11,12^. Besides MIG1, other negative transcriptional regulators, such as ACE1, RCE1, CTF1, etc. have also been identified^13–15^. In the absence of glucose and other fermentable sugars, MIG1 is phosphorylated at a conserved S319 site by protein kinase A, leading to its degradation^16,17^. Degradation of MIG1 marks the arrival of carbon stress, and fungi start responding to the presence of alternate, non-fermentable carbon sources like cellulose^17^.

The induction of cellulose hydrolyzing enzymes is governed at transcriptional level by the positive regulators. These transcriptional activators of cellulase vary among filamentous fungi; XYR1 is the key transcriptional activator in *T. reesei*, and CLR-1 and CLR-2 are set of conserved transcriptional activators in *Neurospora crassa*. In *Aspergillus niger*, XYR1, CLR-2 and to a lesser extent, CLR-1 regulate cellulase expression upon induction. Similar to archetypical GAL4 transcription factor in yeast, CLR-1, CLR-2, and XYR1 are members of the Zn(II)_2_Cys_6_ family of transcription factors having binuclear Zn–Cys binuclear cluster-type DNA-binding domain and activators of genes of secondary carbon sources. Beside these, homologs of ACE3, BGL-R, CRZ1, VIB1, etc., have also been identified as transcriptional activators of lignocellulase in filamentous fungi^15,18–21^.

Lignocellulase production is also influenced by environmental factors like Ca^2+^, light, and pH under carbon stress^15,22,23^. Studies on *T. reesei* indicated that cAMP levels are augmented during carbon stress. This activates phospholipase C, leading to Ca^2+^ burst in the intracellular spaces and positive regulation of key cellulolytic genes^22,23,26^. As a result, calcineurin-dependent calcium signaling upregulates the expression of transcriptional activator XYR1, via calcineurin-responsive zinc finger transcription factor (Crz1), in addition to other unknown factors^25^. Increased expression of XYR1 in turn upregulates cellulolytic gene expression. However, we did not find any report describing the similar effect of calcium signaling on transcriptional activators, CLR-1 and CLR-2.

Additionally, G protein receptors relay cellulase inducing signal by activating Gα subunits, GNA1/3, and elevate cAMP levels to activate Protein kinase A (PKA) pathway in *T. reesei*^27,28^. Another recent study highlighted the role of GNA3 as well as regulator of G-protein signaling, RGS2 in extracellular cellulase production in *N. crassa*^29^. Furthermore, studies on *Aspergillus nidulans* and *N. crassa* identified the activation of the HOG1 MAPK pathway during cellulase production^30,31^. Another study on *N. crassa* indicated high oxygen consumption by fungi to meet increased demands for protein folding in ER during the trafficking of cellulases^32^. Despite the diverse studies, mechanistic understanding of cellulase production post-transcription remained elusive.

The present study was conducted with the aim of identifying the signaling networks governing the extracellular production of lignocellulolytic enzymes in *P. funiculosum* NCIM1228. *P. funiculosum* has recently been identified for its superior hyper-cellulolytic secretome. GH7 cellobiohydrolases (CBH1) is the crucial enzyme for cellulose breakdown in filamentous fungi^33^. *P. funiculosum* CBH1 exhibited an 18-fold higher turnover rate, six-fold higher catalytic efficiency, and 26-fold higher enzyme-inhibitor complex equilibrium dissociation constant (K_i_) than CBH1 of *T. reesei*^34^. Also known for higher β-glucosidase levels, the secretome of *P. funiculosum* is a sustainable substitute for *T. reesei*-based secretomes for wide-ranging industrial applications^12^. A molecular toolbox was created^35^ and the fungus was genetically engineered to replace the functional allele of the catabolite repressor with Mig1 deletion cassette^11^. The resultant strain, ΔMig1 (also called *Pf*Mig1^88^), exhibited catabolite de-repression on most of the enzymes of alternate carbon utilization. Since the key transcriptional activator of cellulase vary among different fungi, we began by identifying the transcription factor responsible for cellulase production in *P. funiculosum* NCIM1228. Advanced transcriptomics identified CLR-2 as the key transcriptional activator of cellulase in *P. funiculosum* in response to induction by the polymeric substrate cellulose. Functional studies of Clr-2 overexpressing mutants revealed its limited role beyond cellulase transcription. We next studied the effect of Ca^2+^-signaling on cellulase transcription, translation, and secretion, and found that the presence of both cellulose and calcium was necessary for cellulase translation and secretion. RNA-seq transcriptomics identified SNF1 AMPK as a Ca^2+^-activated kinase, upregulated in cellulose during carbon stress. Comparative functional analysis and quantitative proteomics of NCIM1228 and ΔSnf1 strains identified the key functions of Ca^2+^-signaling and SNF1 to facilitate cellulase secretion.

## Results

### Increased Ca^2+^ level in the growth medium relieves the post-transcriptional bottleneck in cellulase production

We set out by identifying the key transcription factors (TFs) involved in the regulation of cellulase gene expression in *P. funiculosum*. We analyzed the global transcriptome of log-phase cultures of NCIM1228 and ΔMig1 (the catabolite de-repressed mutant of NCIM1228) grown in glucose and cellulose by RNA-seq (Fig. 1a,b). Comparative analysis of cellulose/glucose detected upregulation of 38 TFs in NCIM1228 and 17 TFs in ΔMig1 in cellulose, while 14 of these were detected in both (Fig. 1c,d). We argued that TFs responsible for cellulase transcriptional activation must be induced in both NCIM1228 and ΔMig1 and should exhibit higher transcript levels in catabolically derepressed ΔMig1. Therefore, 24 TFs exclusive to NCIM1228 and 3 TFs exclusive to ΔMig1 were ignored (Fig. 1d). There were 6 common TFs that were induced in both NCIM1228 and ΔMig1. Among these, 3 TFs had equal transcript levels in both NCIM1228 and ΔMig1 and were identified as homologs of CTF1a (TF for cutinase)^36^, ACU15 (TF for acetate utilization)^37^, and ATF21 (TF for spore maturation)^30^. The other three common TFs demonstrated higher levels in ΔMig1 and were found to be the homologs of CLR-2 (TF for cellulase)^5,19^, UME6 (TF for early meiotic genes)^40^, and KLF1 (TF for G0 phase longevity)^41^, respectively (Fig. 1d). Out of the six shortlisted TFs, CLR-2 and CTF1a seemed relevant to cellulase induction and, therefore, were chosen for further analysis (Fig. 1e). Additionally, we also shortlisted 88_0.112 (CTF1b) for having 33% homology to cutinase regulator of *Fusarium solani*^42^ (Fig. 1e). We re-confirmed the transcript levels of the three shortlisted TFs in glucose and cellulose by real-time PCR and found them in sync with the transcriptomic results (Fig. 1f).

**Fig 1.**
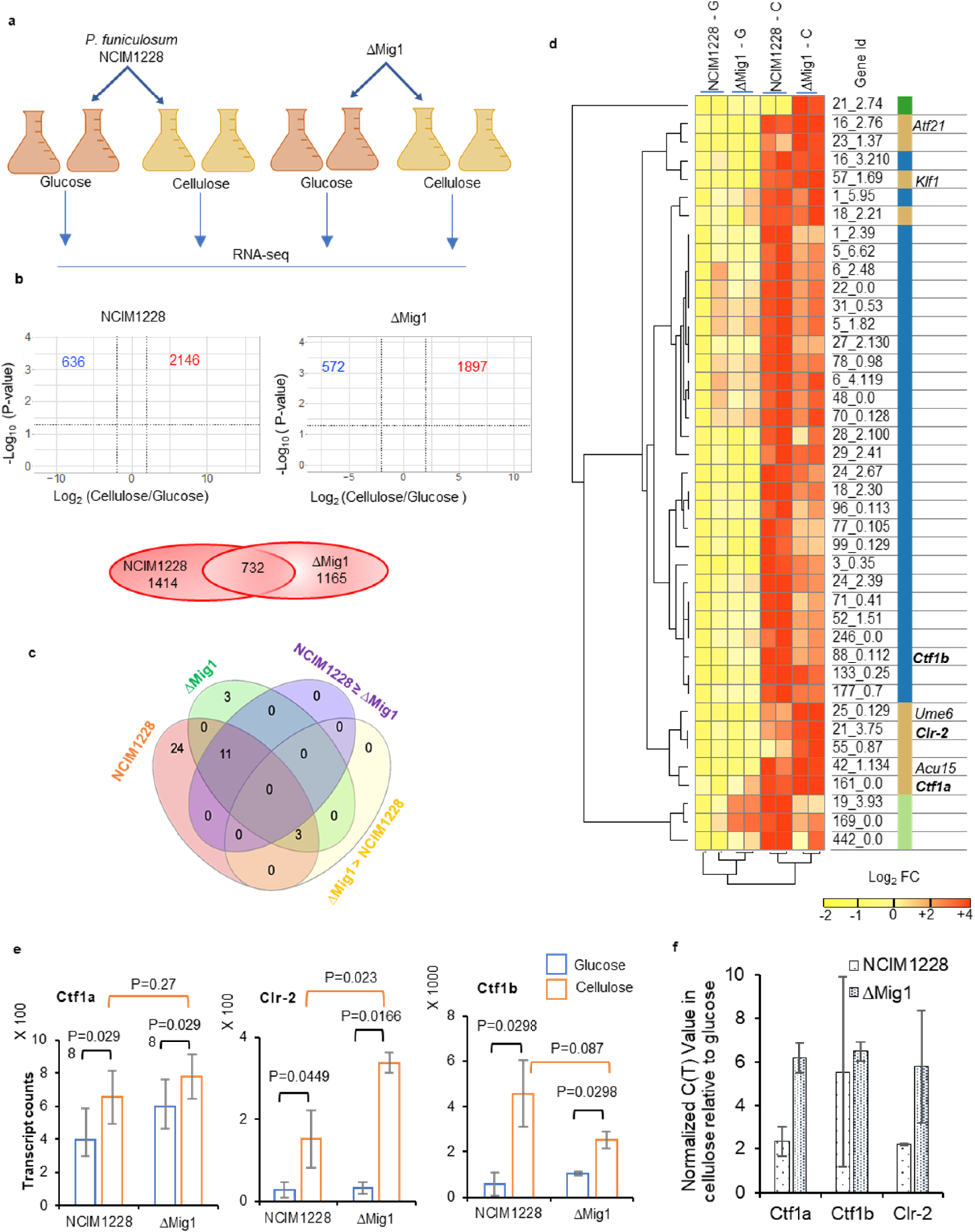
Cellulose induced transcription factors in *P. funiculosum* NCIM1228 and ΔMig1 identified by RNA-seq. **(a)** RNA-seq experimental design to conduct transcriptomic analysis of NCIM1228 and ΔMig1 grown in Mandels medium having 4% glucose (MM glucose) and 4% cellulase (MM cellulose) for 72 hours. **(b)** Volcano plot of differential expressed genes in cellulose respective to glucose in NCIM1228 and ΔMig1. Venn diagram shows the common upregulated genes between NCIM1228 and ΔMig1. **(c)** Venn diagram of common and exclusive transcription factors (TFs) upregulated in NCIM1228 and ΔMig1; 11 of the common TFs showed higher log2 fold change in NCIM1228 than ΔMig1 whereas 3 of them showed an opposite trend. **(d)** Hierarchical and K-means clustering of normalized FPKM abundance (log_2_) of 41 TFs in NCIM1228 and ΔMig1 grown on glucose and cellulose. Four clusters formed are TF exclusively upregulated in ΔMig1 grown on cellulose (1 TF); TFs exclusively upregulated in NCIM1228 (3TFs); TFs upregulated in both NCIM1228 and ΔMig1 with higher FPKM values in NCIM1228 grown on Cellulose (30 TFs); TFs upregulated in both NCIM1228 and ΔMig1 with higher FPKM values in ΔMig1 grown in Cellulose (4 TFs). Three transcription factors related to biomass degradation were identified: CLR-2 (TF for cellulase); CTF1a and CTF1b (TFs for cutinase). **(e)** Transcript counts of CTF1a, CLR-2 and CTF1b in NCIM1228 and ΔMig1 grown in glucose and cellulose Statistical significance was determined by one tailed, unequal variance t-test. **(f)** Quantitative RT-PCR of selected TFs in NCIM1228 and ΔMig1 grown in MM glucose and MM cellulose for 72 hours. C(T) value in cellulose was normalized to that in glucose.

Genes encoding CLR-2, CTF1a, and CTF1b were over-expressed individually in NCIM1228 under their native promoters (Fig. 2a-c, Supplementary Fig. 1), and the increased levels of respective transcripts were confirmed by real-time PCR in over-expressed mutants (Fig. 2d,e). The mutants were grown in cellulosic medium and the secretomes obtained were examined for the total cellulase activity by filter paper unit (FPU) assay (Fig. 2f). Out of the three, P_Clr2_*Clr-2*/NCIM1228 exhibited 2-fold higher FPU levels than NCIM1228; even transcript levels of key cellulase genes, namely, cellobiohydrolase1 (*cbh1*), endoglucanase (*eg-GH45*), β-glucosidase (*bgl-GH3*) and xylanase (*xyl-GH10-CBM1)* were found upregulated in both glucose and cellulose (Fig. 2g). *ctf1a*, and *ctf1b* over-expression didn’t had any effect on cellulase as well as biomass hydrolyzing capacity of the NCIM1228 secretome, however cutinase activity decreased by several fold; the two transcription factors actually behaved as repressors of cutinase under cellulosic conditions (Supplementary Fig. 2).

**Fig 2.**
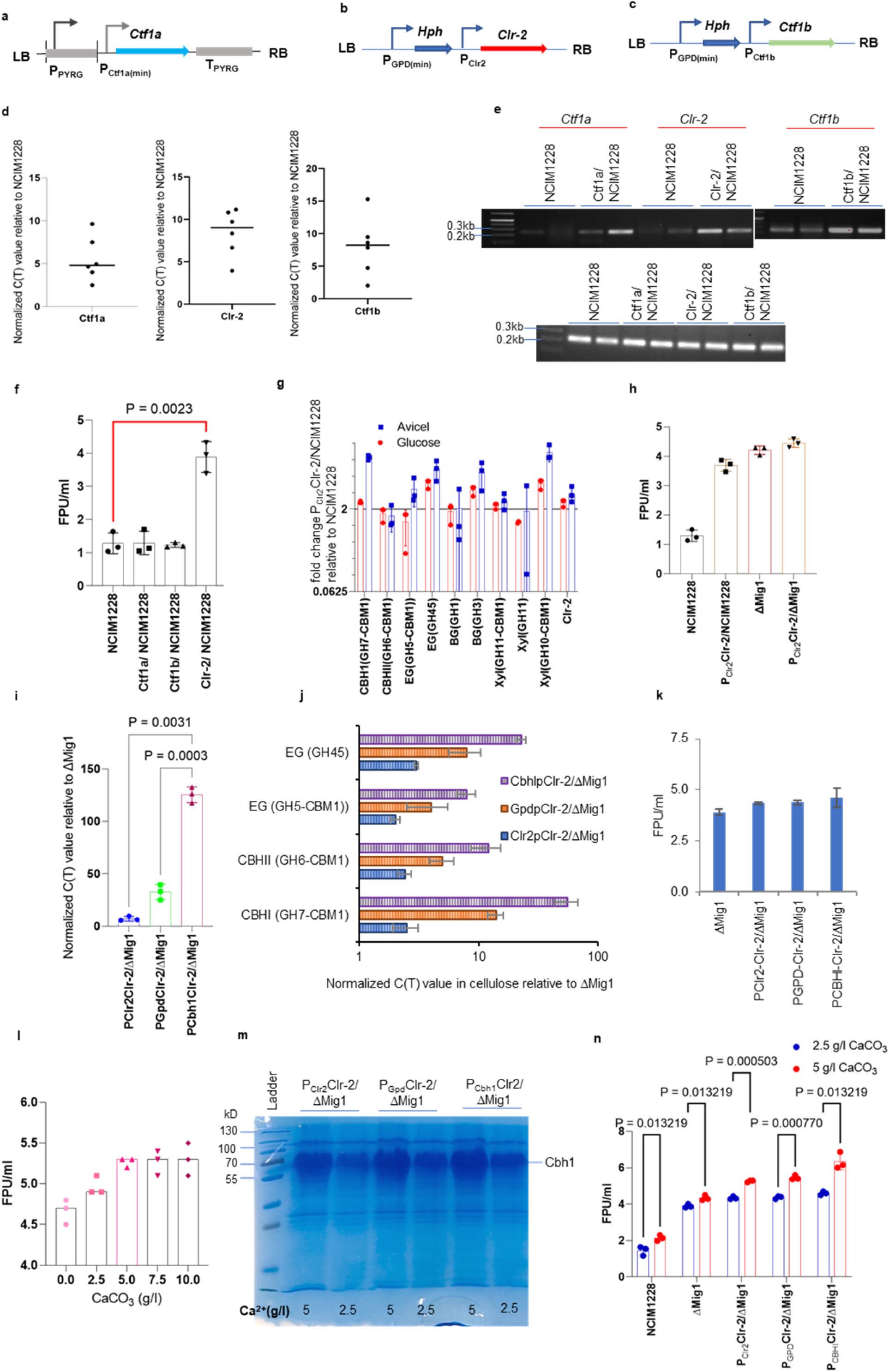
CLR-2 overexpression leads to increased cellulase when complemented with increased Ca^2+^ in the growth medium. Schematic representation of expression cassettes of *ctf1a* **(a)**, *ctf1b* **(b)**, and *clr-2* **(c). (d)** Over expression of *ctf1a* (left), *ctf1b* (middle), and *clr-2* (right) in their respective over-expressed strain measured by real-time PCR in the corresponding transformants. **(e)** Technical replicates of *ctf1a, clr-2* and *ctf1b* PCR products reflecting transcript levels in their respective overexpressed strains. Lower panel shows the technical replicates of tubulin expression in all the strains. **(f)** Total cellulase activity of secretomes measured by FPU assay. **(g)** Transcript levels of cellulases in Clr-2/NCIM1228 relative to NCIM1228, when grown in MM glucose and MM cellulose for 72 hours. **(h)** Total cellulase activity of P_Clr2_Clr-2/ΔMig1 secretome compared with parent strains by FPU assay. RT-PCR showing Transcript levels of **(i)** *clr-2* and **(j)** cellulase genes, when expressed under native, *gpd* (constitutive), and *cbh1* (inducible) promoters. **(k)** Total cellulase activity of secretomes obtained, when *clr-2* overexpressed strains were grown in RCM medium. **(l)** Total cellulase activity of secretomes obtained when P_Clr2_Clr-2/ΔMig1 was grown in RCM medium having varying levels of CaCO_3_. Secretome profile on 12% SDS-PAGE **(m)**, and total cellulase activity **(n)** by FPU assay, of *clr-2* mutants grown in RCM medium having 2.5g and 5.0g CaCO_3_. All the experiments were repeated three times; statistical significance determined by one tailed, unequal variance t-test.

Next, we over-expressed *clr-2* under its native promoter in ΔMig1 strain and, expected an exponential increase in cellulolytic attributes accredited to the reinforced activation mechanisms, conjoint with diminished negative regulation^2,43^ (Supplementary Fig. 3a). However, P_Clr2_*Clr-2*/ΔMig1 secretome showed marginal supremacy over the parent strains (Fig. 2h, Supplementary Fig. 3b-f); similar results were obtained from comparative biomass hydrolysis (Supplementary Fig. 3g,h). Dwelling further into the transcriptomic data made us realize extremely low levels of *clr-2* transcript compared with housekeeping genes. Therefore, we over-expressed *clr-2* under *PfCbh1* inducible promoter (the top-ranked gene expressed in cellulose), and a strong constitutive *gpd* promoter (Supplementary Fig. 1e,j-k). Quantification by real-time PCR found *clr-2* transcript levels in the order P_Cbh1_>P_Gpd_>_PClr-2_ (Fig. 2i); similar pattern was observed in the transcripts for downstream CLR-2 regulated cellulases (Fig. 2j). However, differential expression of *clr-2* transcript and cellulases had minimal effect on secretion levels of the cellulases (Fig. 2k). Bioinformatic analysis of CLR-2 protein predicted Gal4 DNA binding domain (35-80 aa), nuclear localization signal^44^ (45-65 aa) and fungal TF specific domain (355-436 aa). A recent study on CLR-2 suggested the removal of the middle regulatory region of CLR-2 (248-646 aa) to increase cellulase production^45^. In our case, however, a similar strategy resulted in reduced cellulase levels in the ΔMig1 strain; perhaps the removal of the regulatory region from CLR-2 made it non-functional (Supplementary Fig. 4).

Studies on *Trichoderma reesei* indicated the role of Ca^2+^ signaling on hyphal growth and cellulase induction^22,24,25^. In our previous reports, we used 50 mg/l CaCl_2_ or 0.5 g/l CaCO_3_ in the cellulase-inducing RCM medium^22,24,25^. We next reasoned if increasing the Ca^2+^ levels would impact the cellulase production^23,24^. So, we cultured P_Clr2_*Clr-2*/ΔMig1 in RCM medium having 50 mg/l of CaCl_2,_ with varying levels of CaCO_3_ from 0 g/l to 10 g/l; and achieved maximum activity of 5.1 FPU/ml at 5 g/l (Fig. 2l). Next, we cultured all three Clr-2 over-expression mutants in the RCM medium having 5 g/l CaCO_3_ and obtained heightened cellulase levels in the order P_Clr-2_<P_Gpd_<P_Cbh1._ P_Cbh1_*Clr-2*/ΔMig1 secretome had maximum cellulase activity of 6.3 FPU/ml (Fig. 2m,n). Increasing the calcium concentration complemented the increase in *clr-2* transcripts and delivered higher cellulase production in the same order.

### Ca^2+^ regulates cellulase production at the protein level

To understand the role of Ca^2+^ in cellulase production, we first examined if a lack of Ca^2+^ affects hyphal growth in glucose and cellulose. The effect was negligible in both NCIM1228 and ΔMig1 (Fig. 3a). Markedly, even transcriptional induction of *clr-2* as well as cellulases remained unaffected in absence of Ca^2+^ in cellulose medium (Fig. 3b). However, both the strains grown without calcium secreted negligible levels of key cellulases (Fig. 3c). In-gel fluorescent (MUG) assay and Western blotting with anti-CBH1 antibody confirmed reduced β-glucosidase and cellobiohydrolase 1 levels, respectively (Fig. 3c). Quantitative evaluation of the secretomes for four major cellulolytic enzyme classes (exocellulase (Avicelase), endocellulase (CMCase), β-glucosidase (pNPGase), and xylanase) revealed reduced enzyme activities in the order, exocellulase>endocellulase>β-glucosidase>xylanases (Fig. 3d-g). FPU levels fell near to zero levels due to the absence of exocellulase and endocellulase in the secretome (Fig. 3h). Notably, glucoamylase secretion remained unaffected (Fig. 3c,3i).

**Fig 3.**
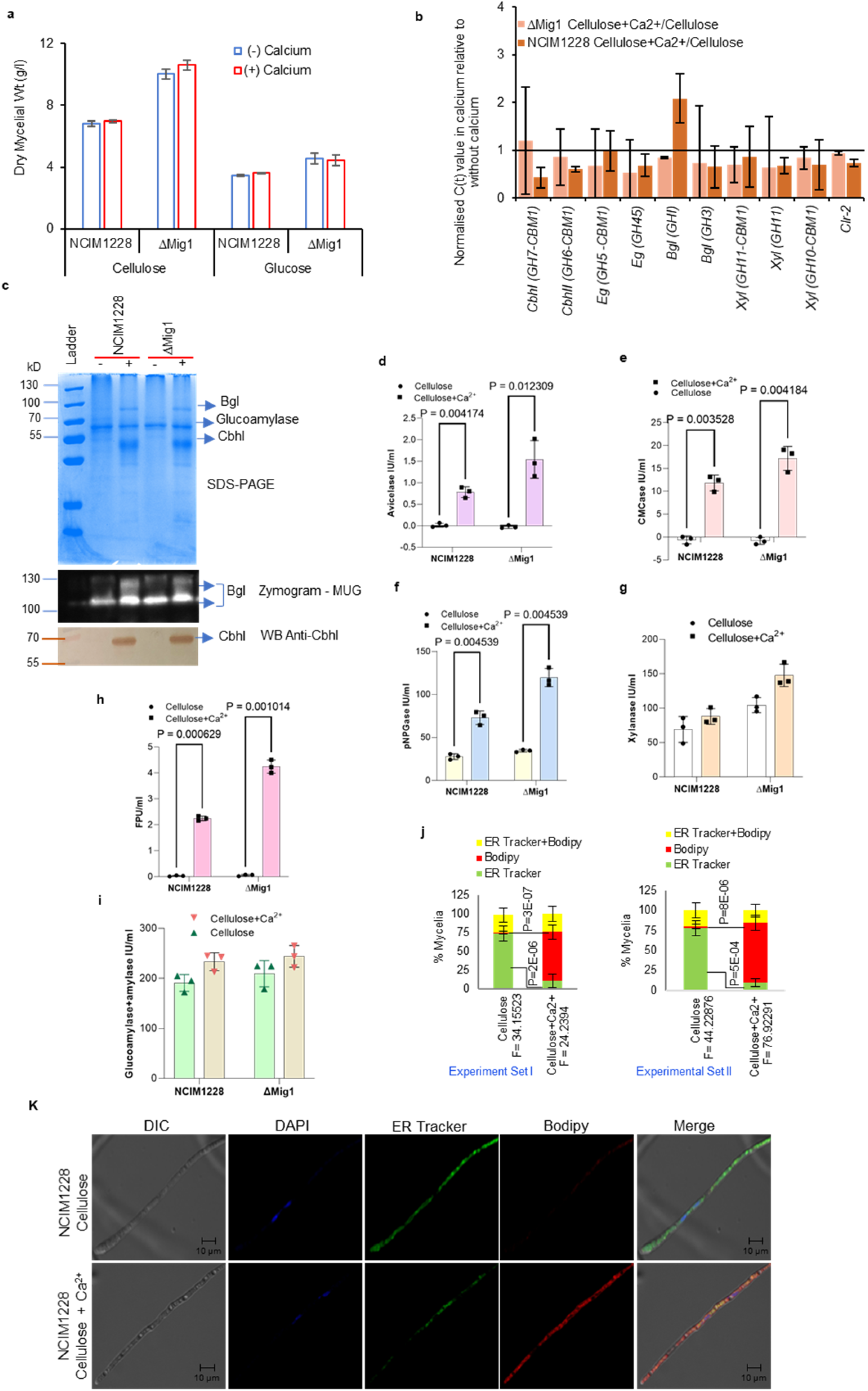
Post-transcriptional events leading to cellulase production are Ca^2+^ dependent. **(a)** Dry mycelial weight of 24-hour cultures of NCIM1228 and ΔMig1 in Mandels medium having 4% glucose and 4% cellulose with and without 50mg/l CaCl_2_. **(b)** Impact of calcium on transcript levels of important cellulases in NCIM1228 and ΔMig1 grown in glucose (30 hours) and cellulose (60 hours) by RT-PCR. **(c)** SDS-PAGE, in-gel MUG β-glucosidase zymogram and Western blot with anti-CBH1 of NCIM1228 and ΔMig1 secretome showing the effect of calcium on total secretome profile, β-glucosidase and cellobiohydrolase levels, respectively. NCIM1228 and ΔMig1 secretomes were also examined for **(d)** Exocellulase (Cellulosease), **(e)** Endocellulase (CMCase), **(f)** β-glucosidase (pNPGase), **(g)** xylanases, **(h)** total cellulase activity by Filter Paper Unit (FPU) assay, and **(i)** amylase activity. All the experiments were repeated three times; statistical significance determined by one tailed, unequal variance t-test. **(j)** percentage NCIM1228 showing staining with ER-tracker and Bodipy when grown in cellulose with and without calcium. The experiment was conducted twice and mycelia (total = 200) in each experiment was counted from 8 slides/condition. All pictures were acquired at fixed laser intensity and exposure in a given experiment. **(k)** Representative confocal microscopy images of NCIM1228 mycelium grown in cellulose with and without calcium and stained with ER tracker, Bodipy, and DAPI for one hour before imaging.

Endoplasmic reticulum (ER) is known to be the storehouse of calcium and more recently, a study in epithelia normal rat kidney (NRK) cells linked calcium signaling to protein trafficking and secretion^46^. Intrigued by the role of Ca^2+^ in the cellulase production, we asked if Ca^2+^ depletion impacts protein production by causing defects in ER and Golgi morphology. Mycelia grown in glucose and glucose+Ca^2+^ had the same phenotype with well stained ER as well as Golgi. However, 75% of the mycelia grown in cellulose without Ca^2+^ showed extended ER-tracker-stained ER with sparse, circular bodipy-stained Golgi (supplementary Fig. 5). On the contrary, >80% of the mycelia grown in cellulose+Ca^2+^ exhibited the reverse pattern with distended Golgi (Fig. 3j,k). The altered pattern of ER and Golgi in cellulose+Ca^2+^ might be due to increased trafficking of proteins from ER to Golgi for secretion. This suggested the role of calcium in the post-transcriptional processes like cellulase translation or secretion.

### SNF1 AMPK is a Ca^2+^-activated kinase during carbon stress

Calcium signal is generally relayed by Ca^2+^-binding sensory proteins such as Calmodulin, calcium dependent protein kinases and calcineurin etc. We therefore mined NCIM1228 and ΔMig1 transcriptomic data for Ca^2+^-activated kinases, and found six of them being differentially regulated in glucose and cellulose (Fig. 4a). Among the six, the genes encoding two kinases, 87_0.23 (SNF1, AMP activated Kinase (AMPK)) and 75_0.37 (SSP1, Calcium activated kinase kinase (CaMKK)) were upregulated in both NCIM1228 and ΔMig1 in cellulose, with higher fold change in ΔMig1. Literature search also indicated the role of SNF1 in transcriptional de-repression of galactose utilization genes in yeast^47–50^. Besides, SNF1 inhibited amino acid biosynthesis by mTOR inactivation, and activated TFs favoring gluconeogenesis and alternate carbon utilization in *Saccharomyces cerevisiae*^48,49^. SNF1 homolog in *P. funiculosum* is a 823aa protein with conserved serine-threonine kinase domain (87-420aa), which exhibits 67%, 46% and 44% sequence identity to SNF1 protein from *Aspergillus niger, S. cerevisiae* and *Neurospora crassa*, respectively. In *S. cerevisiae*, SNF1 gets phosphorylated in the presence of alternate carbon sources under low-glucose levels^49,51,52^. We found the similar pattern in *P. funiculosum* by Western blotting with anti-phospho-SNF1 (P-SNF1); SNF1 remains unphosphorylated in glucose and gets phosphorylated in cellulose and other secondary carbon sources, only in the presence of Ca^2+^ (Fig. 4b,c).

**Fig 4.**
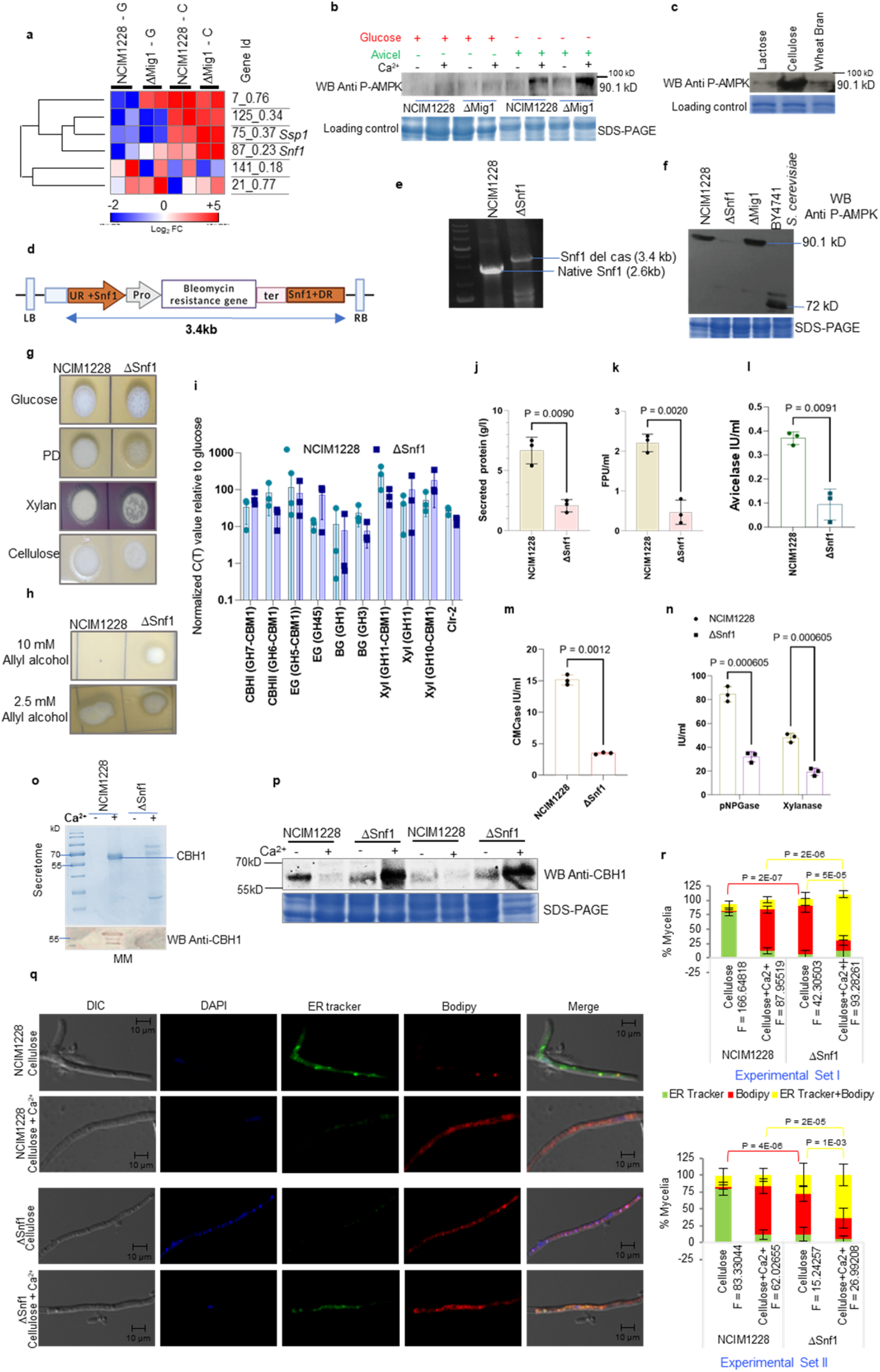
Regulation of cellulase production by Snf1 AMPK. **(a)** Heat map of differentially regulated Ca^2+^-activated kinases identified in NCIM1228 and ΔMig1. **(b)** NCIM1228 was grown in MM glucose and MM cellulose with and without Ca^2+^; phosphorylation status of Snf1 in whole cell extracts was examined by Western blot with Anti-phospho-Snf1 AMPK. **(c)** Anti P-AMPK Western blot with NCIM1228 mycelial extract, when grown in MM lactose, MM cellulose and MM wheat bran (hemicellulose). **(d)** Schematic of *snf1* disruption cassette transformed into NCIM1228. **(e)** PCR with Snf1 Xho1F and Snf1 MauB1R primers, confirming replacement of 2.6kb native copy with that of 3.4kb disruption cassette. **(f)** Anti P-AMPK Western blot with NCIM1228, ΔSnf1 and ΔMig1 mycelial extract, when grown in MM cellulose; *S. cerevisiae* BY4741 grown in lactose was taken as control for P-AMPK. **(g)** Equal number of spores of NCIM1228 and ΔSnf1 were spotted on SC growth medium having 2% glucose, potato dextrose, xylan, cellulose, and **(h)** 25mM Allyl Alcohol (AA). **(i)** Log phase cultures of NCIM1228 and ΔSnf1 were grown in MM cellulose for 72 hours. RT-PCR showing the effect of *snf1* deletion on transcript levels of cellulases. Both NCIM1228 and ΔSnf11 were grown in cellulosic RCM growth medium for five days. NCIM1228 and ΔSnf1 secretomes were examined for **(j)**, total protein content of the secretome **(k)** total cellulase activity by FPU assay, and individual classes of enzymes namely - **(l)** exocellulase (Avicelase), **(m)** endocellulase (CMCase) **(n)**, β-glucosidase (PNPGase) as well as xylanases. All the experiments were repeated three times in triplicates; statistical significance was determined by one tailed, unequal variance t-test. NCIM1228 and ΔSnf1 were grown in MM cellulose for 72 hours. 10 µl of secretomes were collected and 25µg of total mycelial proteins were electrophoresed on 12% SDS-PAGE. **(o)** Secretomes and **(p)** total mycelial extracts were examined for cellobiohydrolase I (CBHI) by Western blotting with Anti-CBHI. for total cellulase activity **(q)** Representative confocal microscopy images of NCIM1228 and ΔSnf1 mycelium grown in cellulose with and without calcium and stained with ER tracker, Bodipy, and DAPI for one hour before imaging. **(r)** Percentage NCIM1228 and ΔSnf1 mycelia showing staining with ER-tracker and Bodipy when grown in cellulose with and without calcium. The experiment was conducted twice and mycelia (total = 200) in each experiment was counted from 8 slides/condition. All pictures were acquired at fixed laser intensity and exposure in a given experiment.

### SNF1 AMPK is an effector molecule that facilitates the cellulase secretion in response to calcium

We next disrupted *snf1* gene at its kinase domain; a 597bp coding region of *snf1* was replaced with bleomycin resistance cassette, the resultant disruption cassette was cloned in a binary vector and transformed into NCIM1228 by *Agrobacterium*-mediated method (Fig. 4d). *snf1* disruption was confirmed by PCR showing the original gene replacement by disruption cassette (Fig. 4e). We further confirmed *snf1* deletion by anti-P-SNF1 Western blot, in which ΔSnf1 showed no band in cellulose (Fig. 4f). As compared to NCIM1228, ΔSnf1 showed sluggish growth on glucose, potato dextrose, xylan as well as cellulose, but exhibited increased resistance towards allyl alcohol (Fig. 4g,h); this might be due to the reduced production of enzymes of secondary carbon utilization and alcohol dehydrogenase. Yet, transcriptional response to cellulose was intact in ΔSnf1 (Fig. 4i). Further, ΔSnf1 secretome displayed abated protein content even in the presence of Ca^2+^; diminished protein and cellulase contents were confirmed by quantification using BCA method and FPU assay, respectively (Fig. 4j,k). Among different classes of cellulases, exocellulase and endocellulase levels were highly affected followed by β-glucosidase and xylanases (Fig. 4l-n). This clearly indicated the role of Ca^2+^-activated SNF1 AMPK in the extracellular production of cellulolytic enzymes. A recent study indicated the upregulation of several genes encoding ER chaperones along with cellulases after Ca^2+^ addition in *T. reesei*^25^. Based on this, We next questioned if the Ca^2+^-activated SNF1 AMPK exerts its control at cellulase translation and maturation level. For this, we detected extracellular as well as intra-mycelial CBH1 level in 72-hour cultures of NCIM1228 and ΔSnf1 grown in cellulose with and without calcium by Western blotting with anti-CBH1 (Fig.4o,p). The secretome of ΔSnf1 didn’t show any band corresponding to CBH1 on SDS-PAGE gel or Western blotting with anti-CBH1 antibody (Fig.4o). We detected diminutive intra-mycelial CBH1 levels in NCIM1228 and ΔSnf1 grown only on cellulose, negligible in NCIM1228 grown in cellulose+Ca^2+^, but elevated levels of CBH1 in ΔSnf1 grown in cellulose+Ca^2+^ (Fig. 4p). Combining intra-mycelial CBH1 levels with those from secretome (Fig. 4o,p), we could infer that the Ca^2+^ is necessary for both translation and secretion of cellulase and the Ca^2+^-activated SNF1 AMPK in an effector molecule that facilitates the cellulase secretion. We made this observation because we detected diminutive levels of CBH1 both intracellularly and extracellularly in both NCIM1228 and ΔSnf1 grown without calcium, which indicates that there is a very little translation of cellulase in the absence of calcium. In the presence of both cellulose and calcium, we detected elevated levels of CBH1 in NCIM1228 secretome and diminutive CBH1 levels inside the mycelia, indicating continuous secretion of CBH1 upon induction. ΔSnf1 showed similar phenotype as NCIM1228 when grown without calcium. However, we found elevated levels of CBH1 inside the ΔSnf1 mycelia grown on cellulose+Ca^2+^ and negligible CBH1 extracellularly confirms intact cellulase translation mechanisms but disrupted secretion in ΔSnf1 (Fig.4o,p). By this experiment, we could conclude that induction of cellulase translation is activated by Ca^2+^ signaling independent of SNF1. The results also suggest that the increased cellulase production observed with increase in Ca^2+^ supplementation, might be the effect of increased cellulase translation, while secretion of cellulases is regulated by Ca^2+^-activated SNF1 AMPK.

We also examined the effect of *snf1* deletion on the ER and Golgi by confocal microscopy (Fig.4q,r); NCIM1228 and ΔSnf1 mycelia were grown in cellulose and cellulose+Ca^2+^ for 72 hours, and stained with ER tracker (ER) and Bodipy (Golgi). All images were captured at fixed laser intensity and exposure time. 75% of the NCIM1228 mycelia grown in cellulose showed well-developed ER, with less than 10% having both ER and Golgi. However, upon the addition of calcium, we observed just the opposite (Fig.4q,r). >70% of NCIM1228 mycelia grown in cellulose+Ca^2+^ had high Golgi content, with ∼10% of the mycelia showing high ER content. On the contrary, >70% of the ΔSnf1 mycelia grown in cellulose showed poorly developed ER-tracker stained ER and increased pattern of circular bodipy-stained Golgi; and in cellulose+Ca^2+^, >65% of the mycelia had both Golgi and ER fully developed and Golgi showed distended morphology as in NCIM1228 (Fig. 4q,r). These results indicated that Ca^2+^ signaling might be inducing cellulase secretion by inciting ER to Golgi transport, strengthening/upregulating the Golgi apparatus and secretory machinery of the mycelia. NCIM1228 grown in cellulose+Ca^2+^ and ΔSnf1 grown in cellulose showing similar ER-Golgi pattern indicates deregulation of the ER to Golgi transport in the absence of SNF1 AMPK. The addition of calcium to ΔSnf1 resulting in similar levels of ER and Golgi fluorescence further strengthens the hypothesis of deregulated production and expression of ER and Golgi proteins in the absence of Snf1 AMPK.

### P-SNF1 AMPK determines upper threshold for Ca^2+^-induced cellulase production

We found above the vital role of Ca^2+^ signaling and Ca^2+^-activated Snf1 AMPK in the cellulase post-transcriptional and secretion process, respectively (Fig. 5a). To gather further insight, we built proteomic profiles of NCIM1228 and ΔSnf1 grown in 4% cellulose with and without calcium (Supplementary Fig. 5). For relative quantification, we added 1µg ^13^C and ^15^C SILu™ APOA-1 (Sigma) to 50 µg protein sample before processing for LC-MS/MS. The whole proteome data obtained for a sample was normalized to the factorial ApoA-1 abundance in the given sample relative to the total ApoA-1 abundance in all the samples. A two-way Anova test identified significantly different proteome maps of NCIM1228 and ΔSnf1 grown in cellulose and cellulose+Ca^2+^ (p<0.05), respectively (Fig. 5b). In cellulose+Ca^2+^ proteome, most of the detected proteins showed much higher abundance than cellulose proteome in both the strains, but the induction was significantly higher in ΔSnf1 as compared to NCIM1228 (Fig. 5c). This was also reflected in the principal component analysis where the proteome profiles grown in cellulose didn’t show a significant difference between the two strains; however, cellulose+Ca^2+^ proteome profiles of NCIM1228 and ΔSnf1 formed separate groups (Fig. 5d). We found 65 proteins upregulated in NCIM1228 and 586 proteins upregulated in ΔSnf1 (Fig. 5e,f); this included 25 (p<0.05) proteins common to both subsets (Fig. 5g). Majority proteins including the common subset exhibited higher abundance levels in ΔSnf1 (Fig. 5g). The common pathways identified in NCIM1228 and ΔSnf1 are central carbon metabolic pathways (TCA cycle, gluconeogenesis, and pentose phosphate pathway), oxidative phosphorylation, proteasomal protein degradation/autophagy, amino acid metabolism, lipid/ vesicle biosynthesis, secretory system including ER transport and processing. As illustrated in Supplementary Fig. 7, the number of proteins representing these pathways are greater in ΔSnf1. Collectively, these common pathways are involved in energy generation during carbon stress, cellulase production and transport machinery. 29 of these upregulated proteins formed an interactome in the evolutionary close *Penicillium marneffei*, as detected STRING database (Fig. 5h).

**Fig 5.**
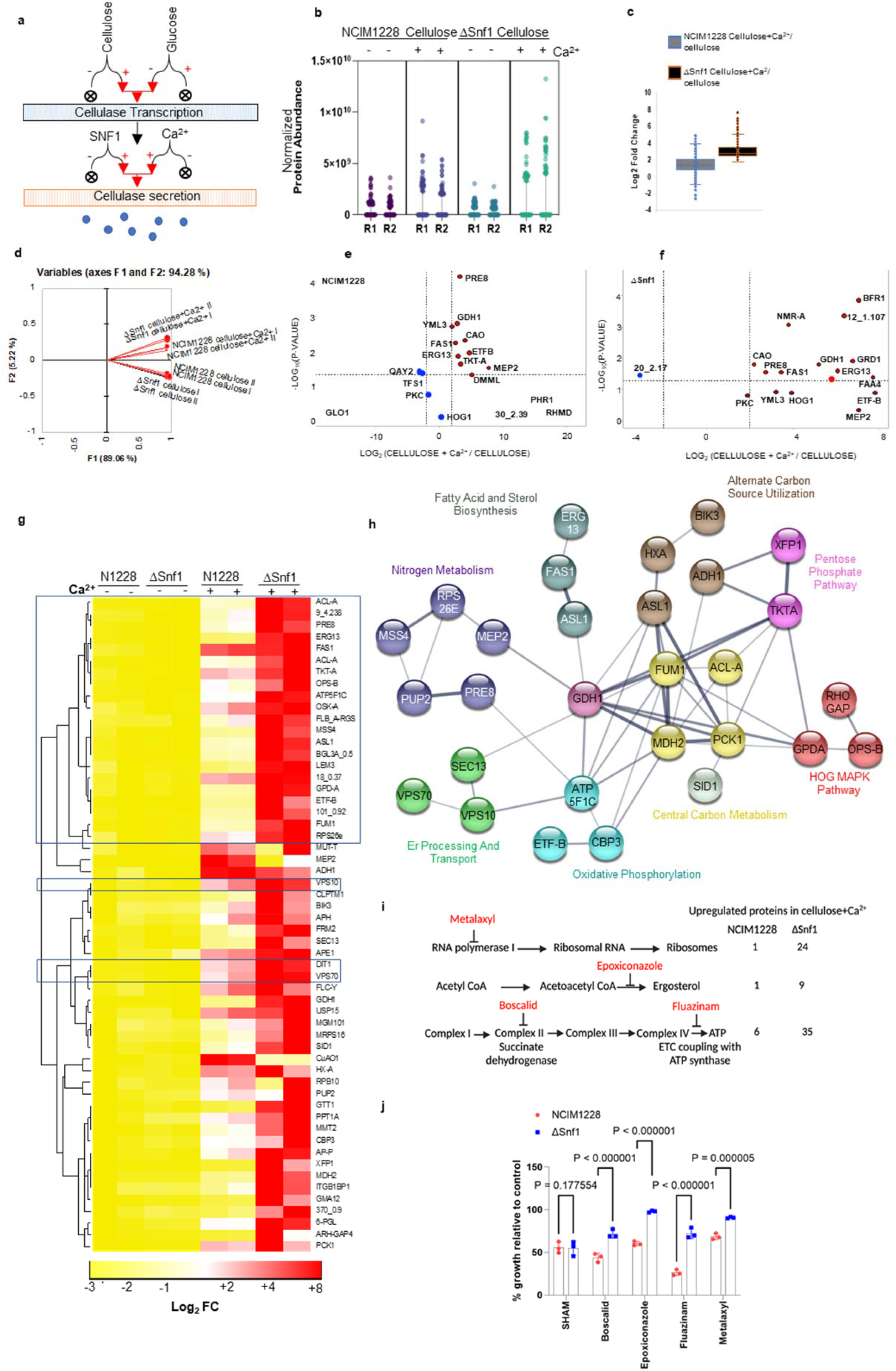
SNF1 AMPK regulates Ca^2+^ activated processes during carbon stress. **(a)** Schematic showing the regulation of cellulase production by different factors; both Snf1 and Ca^2+^ regulate cellulase production post-transcription, we aimed to sought the mechanistic by quantitative proteomics approach. NCIM1228 and ΔSnf1 were grown in MM cellulose with and without calcium for 72 hours and whole mycelial proteins were extracted. 1µg of labelled Apo-1 was added to 50ug of each sample before sample preparation for LC-MS/MS. The proteome data were analyzed by proteome discoverer and normalized to labelled Apo-1. **(b)** Box plot comparing the distributions of data among the replicates as well as different groups. The line represents 5^th^ and 95^th^ whisker boundaries and outliers are represented by dots. A two way ANOVA test without interaction predicted significant difference (F(row)=12.39 with p-value<0.0001; F(column)=687.5 with p-value<0.0001) within the samples. **(c)** Principal component analysis (PCA) showed differential regulation of NCIM1228 and ΔSnf1 grown in cellulose+Ca^2+^. Volcano plot of normalized data of **(d)** NCIM1228, **(e)** ΔSnf1 cellulose+Ca^2+^ proteome. **c**, Normalized protein abundance data of all samples, the lines indicate where 90% of the proteins fall and the dots represent the outliers, within the sample. **(f)** Box plot to show the fold change between cellulose+Ca^2+^/cellulose proteome in NCIM1228 and ΔSnf1. The box represents the 25^th^ percentile and 75^th^ percentile of the data and a line is drawn inside the box at the median (the 50th percentile). **(g)** Heat map of proteins upregulated in both NCIM1228, the boxed proteins form the common subset. **(h)** Important pathways are represented by proteins and interactions being represented by some of the NCIM1228 and ΔSnf1 in cellulose+Ca^2+^ using reference proteins of *Penicillium marneffei* in the String database. **(i)** Schematic of the mode of action of antifungals used in the study and the upregulated proteins in NCIM1228 and ΔSnf1 representing the targeted pathway. **(j)** Antifungal susceptibility of NCIM1228 and ΔSnf1 towards salicylhydroxamic acid (SHAM, Alternate oxidase inhibitor, used as control; EC_50_ 100μg/ml), Boscalid (succinate dehydrogenase (complex II) inhibitor; EC_50_ 1μg/ml), Epoxiconazole (sterol biosynthesis inhibitor; EC_50_ 1.2μg/ml), Fluazinam (uncoupler of oxidative phosphorylation; EC_30_ 0.3μg/ml), and metalaxyl (RNA polymerase I inhibitor; EC_50_ 60μg/ml). The graph represents the % growth (dry mycelial weight) achieved in the presence of antifungals compared with control, after 24h hours of incubation. All the experiments were repeated three times in triplicates; statistical significance was determined by one tailed, unequal variance t-test.

Proteomic studies suggested the down-regulation of cellulose+Ca^2+-^activated processes by Snf1. We verified proteomic results by chemical-genetic approach and determined ΔSnf1 resistance towards antifungals targeting Ca^2+^-activated processes (sterol biosynthesis, mitochondrial respiration and RNA polymerase1)^53–55^(Fig. 5i). At half maximal effective concentration (EC_50_) of these antifungals for NCIM1228, ΔSnf1 showed better growth dynamics than NCIM1228 on Boscalid, Fluazinam, Epoxiconazole, and Metalaxyl. This confirms the upregulation of chemically targeted pathways in ΔSnf1 (Fig. 5j).

Additionally, whole cell proteome of ΔSnf1 cellulose+Ca^2+^ showed increased abundance of CBH1, BGL-GH3, and Xyl(GH10-CBM1) among other CAZymes, validating our earlier discussed findings of zilch effect of Snf1 deletion on cellulase translation. It also confirmed the role of Snf1 in the secretion of cellulases (Fig. 6a). To confirm the abundance of CAZyme RNAs, NCIM1228 and ΔSnf1 were grown in cellulose and cellulose+Ca^2+^ for 72 hours and the transcript levels of *Cbh1, Bgl-GH3*, and *Xyl* (GH10-CBM1) genes were examined in cellulose+Ca^2+^ relative to cellulose. Interestingly, we found much higher fold change of CAZyme transcripts in ΔSnf1 as compared to NCIM1228 (Fig. 6b), which indicated that SNF1 negatively regulates cellulase transcription. While these Ca^2+^-activated pathways and cellulase transcription were also found upregulated in NCIM1228, but to a lesser extent; the SNF1 AMPK driven inhibition is not complete, rather sets the upper threshold levels for regulated cellulase production beyond transcription.

**Fig 6.**
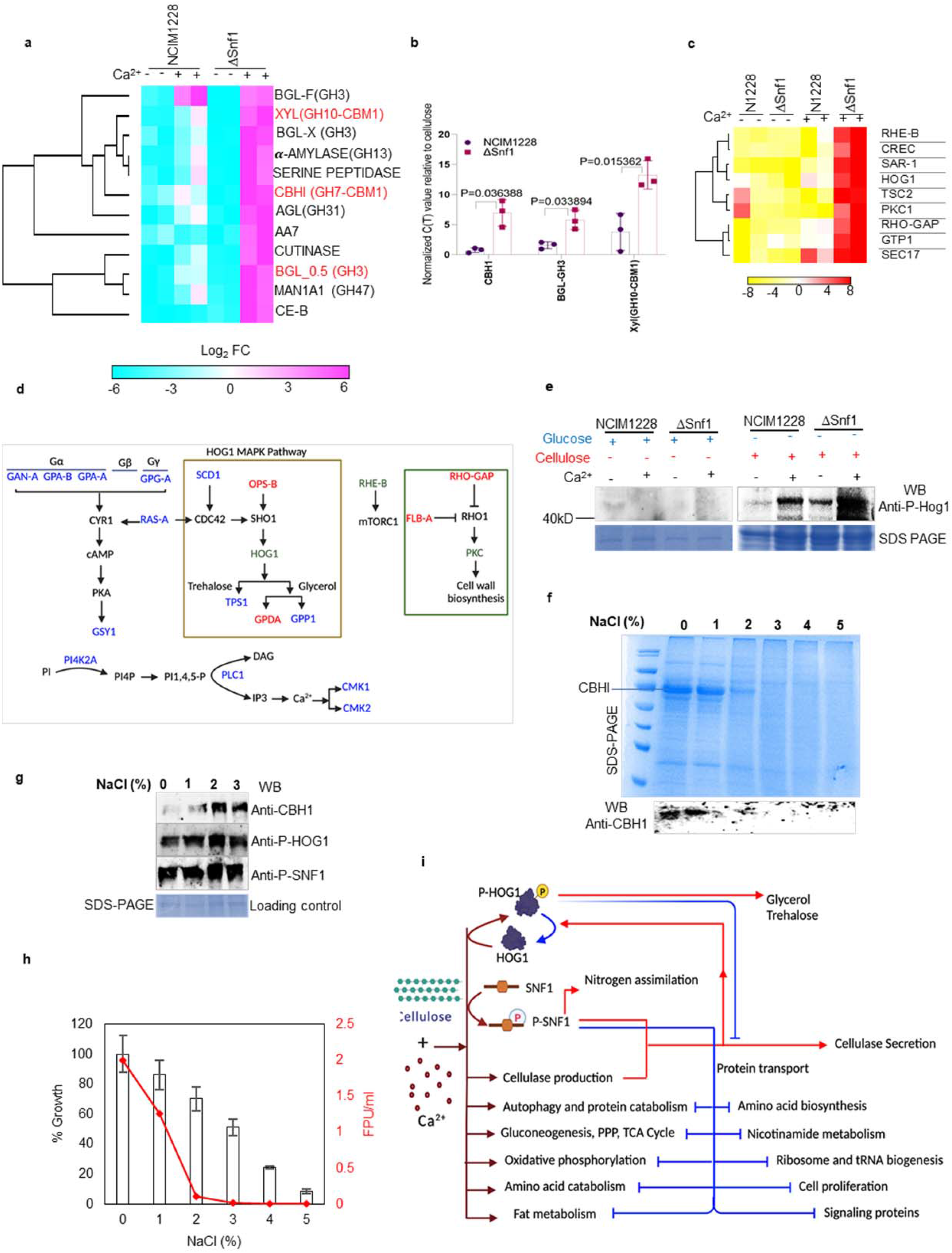
SNF1 AMPK facilitates cellulase secretion by regulating phosphorylation dynamics of Hog1 MAPK. **(a)** Heatmap showing the CAZYmes abundance in NCIM1228 and ΔSnf1 grown in cellulose and cellulose+Ca^2+^. **(b)** Transcript levels of *cbh1, bgl-GH3*, and *xyl(GH10-CBM1)* in NCIM1228 and ΔSnf1 in cellulose+Ca^2+^ relative to cellulose, measured by real-time PCR. **(c)** Heatmap showing protein levels of signaling proteins in the proteomes of NCIM1228 and ΔSnf1. **(d)** Schematic showing upregulated signaling proteins, proteins shown in red were upregulated in both NCIM1228 and ΔSnf1 cellulose+CA^2+^ proteome with respect to their cellulose proteome, those shown in blue were exclusively upregulated in ΔSnf1 cellulose+Ca^2+^ /cellulose proteome, and the ones shown in green were more abundant in ΔSnf1 cellulose+Ca^2+^ as compared to NCIM1228 cellulose+CA^2+^ proteome. **(e)** WB showing P-Hog1 levels in mycelial extracts of NCIM1228 and ΔSnf1 grown in glucose and cellulose with and without calcium. **(f)** SDS-PAGE showing secretome of NCIM1228 grown in RCM medium having varying concentrations of NaCl. **(g)** Graph showing the effect of NaCl on growth and cellulase secretion. The Red lines denote the total cellulolytic capacity of the secretomes of NCIM1228 grown in RCM medium having varying concentrations of NaCl. Black lines denote the percentage growth in 4% glucose achieved at different NaCl concentrations relative to the control having no salt. **(h)** Anti-CBH1, anti-P-Hog1, and anti-P-Snf1 western blots of mycelial extracts of NCIM1228 grown in RCM cellulosic medium supplemented with 1 to 5%NaCl **(i)** Schematic overview of signaling events regulated by cellulose+Ca^2+^ signaling, SNF1 AMPK and HOG1 MAPK in *Penicillium funiculosum* NCIM1228 during carbon stress. Cellulose+Ca^2+^ signaling initiates a series of events involved in deriving amino acid pools from non-carbohydrate sources by means of autophagy and protein catabolism. The amino acid pool is used to produce cellulases as well as meet copious energy needs, for cellulase production, by enhancing amino acid catabolism, gluconeogenesis, TCA cycle as well as mitochondrial respiratory machinery. Ca^2+^ signaling also hyperactivates HOG1 MAPK to combat oxidative stress caused by metabolic bursts. However, hyperactivation of HOG1 MAPK impairs vesicle trafficking causing secretion block. SNF1 AMPK activated by cellulose+Ca^2+^ signaling not only adjusts HOG1 phosphorylation dynamics to remove secretion block, but also regulates cellular processes causing a metabolic burst. Additionally, SNF1 AMPK inhibits glucose-activated pathways like amino acid biosynthesis, ribosome and tRNA biogenesis, cell proliferation and signaling proteins involved.

Besides the common pathways, proteome of ΔSnf1 grown in cellulose+Ca^2+^ exclusively showed upregulation of processes generally activated by mTORC1 complex in response to glucose availability (Supplementary Fig. 8a,b). Most notable ones were amino acid biosynthesis, ribosome and tRNA biogenesis among other anabolic pathways (Supplementary Fig. 8a). This suggests stringent inhibition of mTORC-activated pathways by Ca^2+^-activated P-SNF1 in NCIM1228. Since SNF1 AMPK is known to downregulate mTORC1 driven processes upon the arrival of stress in yeast as well as mammals, these results confirmed fitness of our proteomic analysis.

We also identified the functions of unphosphorylated SNF1 AMPK by comparing NCIM1228 cellulose proteome to that of ΔSnf1 and discovered the down-regulation of stress-response related proteins (HMA5 (copper transporting ATPase), transcription factors (DEF1, BDF1), and nuclear transport regulators (NTF2, SNX12) in ΔSnf1 cellulose proteome (Supplementary Fig. 8c). This suggests that response of SNF1 AMPK to the carbon stress is distinct in the absence of calcium.

### SNF1 AMPK relieves secretion block by regulating HOG1 phosphorylation

We next scrutinized NCIM1228 and ΔSnf1 proteomes for upregulated signaling proteins (Fig. 6c). NCIM1228 cellulose+Ca^2+^ proteome signaling profile hinted at the RHO1 inhibition, and HOG1 MAPK activation *via* SHO1 branch during cellulase production (Fig. 6c,d). Among the upregulated signaling proteins in ΔSnf1, we found increased levels of RheB which is known to activate mTORC1 and gets inhibited by P-SNF1 *via* TSC complex (Fig. 6c,d). Furthermore, ΔSnf1 grown in cellulose+Ca^2+^ showed >2-fold higher abundance of proteins related to conserved cAMP-PKA and Ca^2+^ signaling pathways than those found in NCIM1228, (Fig. 6c,d). Compared with NCIM1228 cellulose+Ca^2+^, HOG1 exhibited >2-fold higher abundance in ΔSnf1 cellulose+Ca^2+^ proteome (Fig. 6c, d). Moreover, we also found two activating proteins RAS-A and SCD1 of SHO1 branch in ΔSnf1, suggesting hyperactivation of HOG1 in ΔSnf1^56–59^ (Fig. 6c,d). Previous reports on *Aspergillus nidulans* and *N. crassa* have shown HOG1 phosphorylation via CDC42-SHO1 branch on non-glucose carbon sources^30,31^. *P. funiculosum* HOG1 is a highly conserved 40.8 kD MAP kinase exhibiting high homology to HOG1 from *P. marneffei* (99.44% identity, 100% coverage), *N. crassa* (92.22% identity, 97% coverage), *T. reesei* (94.19% identity, 96% coverage), and *S. cerevisiae* (84% identity, 96% coverage). We used anti-phospho-p38 antibody to specifically detect phosphorylated HOG1 (P-HOG1) by Western blotting. We found HOG1 phosphorylation in cellulose to be Ca^2+^ dependent in NCIM1228 (Fig. 6e). Further, ΔSnf1 exhibited high P-HOG1 levels even in cellulose, which increased manifold when the cellulosic medium was supplemented with calcium (Fig. 6e). Ca^2+^-activated SNF1, therefore, down-regulates P-HOG1 levels during cellulase production. Our earlier reports in *S. cerevisiae* and *Candida albicans* have shown impeded vesicular and endosomal trafficking by P-HOG1^60^. Salt stress activates HOG1 MAPK in all yeasts and filamentous fungi; we next asked if the fluctuations in P-HOG1 levels caused by salt stress affects cellulase secretion in cellulose (Fig. 6f,g). At 2% and 3% NaCl in glucose, NCIM1228 showed >50% growth; similar conditions in cellulose had no effect on CBH1 translation, but completely abolished cellulase secretion (Fig. 6f-h). As expected, addition of salt increased P-HOG1 and P-SNF1 levels in the cellulose+Ca^2+^ growth medium. This confirms the negative regulation of cellulase secretion by P-HOG1, which is counteracted by SNF1 during cellulase production. We summarize our results in Fig. 6i.

### SSP1 CaMKK phosphorylates SNF1 AMPK

Ca^2+^-activated SNF1 AMPK positively regulates cellulase secretion. This implied that reinforcing the AMPK pathway should result in increased extracellular cellulase production. We found the gene ID 75_0.37 encoding for the most significant CaMKK (homologous to *S. cerevisiae* SSP1 protein with 60% identity) with ∼5-fold higher transcript levels in cellulose than glucose in ΔMig1 in RNA-seq data (Fig. 4a). Real-time PCR of *ssp1* in NCIM1228 and ΔMig1 detected extremely low transcript levels in glucose compared to tubulin, which increased by 3 to 6-fold in cellulose (Fig. 7a). Earlier reports in *Schizosaccharomyces pombe* revealed that SSP1 CaMKK phosphorylates SSP2 AMPK in low glucose conditions^61–63^. Based on this, *ssp1* gene of *P. funiculosum* encoding a 138 kD protein was cloned under the native glucoamylase promoter in a binary vector and transformed into NCIM1228 and ΔMig1 (Fig. 7b). Transformants were confirmed for having an additional copy of *ssp1* by PCR and the increased transcript levels of *ssp1* in glucose and cellulose were confirmed by real-time PCR (Fig. 7c-e). SDS-PAGE profile and FPU assay of secretomes reflected no change in cellulase production by SSP1/NCIM1228 as compared to NCIM1228 (Fig. 7f,g), but SSP1/ΔMig1 secretomes exhibited 40% higher FPU levels than ΔMig1 (Fig. 7f,g). We didn’t find noticeable change between the cellulase transcript levels of SSP1/ΔMig1 and ΔMig1, suggesting the reason of increased activity might be at protein level (Fig. 7h). Since higher transcript levels could also be achieved by *ssp1* transformants on glucose (which is otherwise inhibitory) due to its expression under the control of glucoamylase promoter, we examined if SSP1 could phosphorylate SNF1. Western blotting with anti-P-AMPK found that SNF1 remained unphosphorylated in glucose in both NCIM1228 and ΔMig1, and the state didn’t change in SSP1/NCIM1228 transformants having intact repression mechanisms. However, P-SNF1 levels increased appreciably with SSP1 over-expression in ΔMig1 having disrupted repression mechanisms (Fig. 7i), which could be the reason of increased cellulase production by Ssp1/ΔMig1.

**Fig 7.**
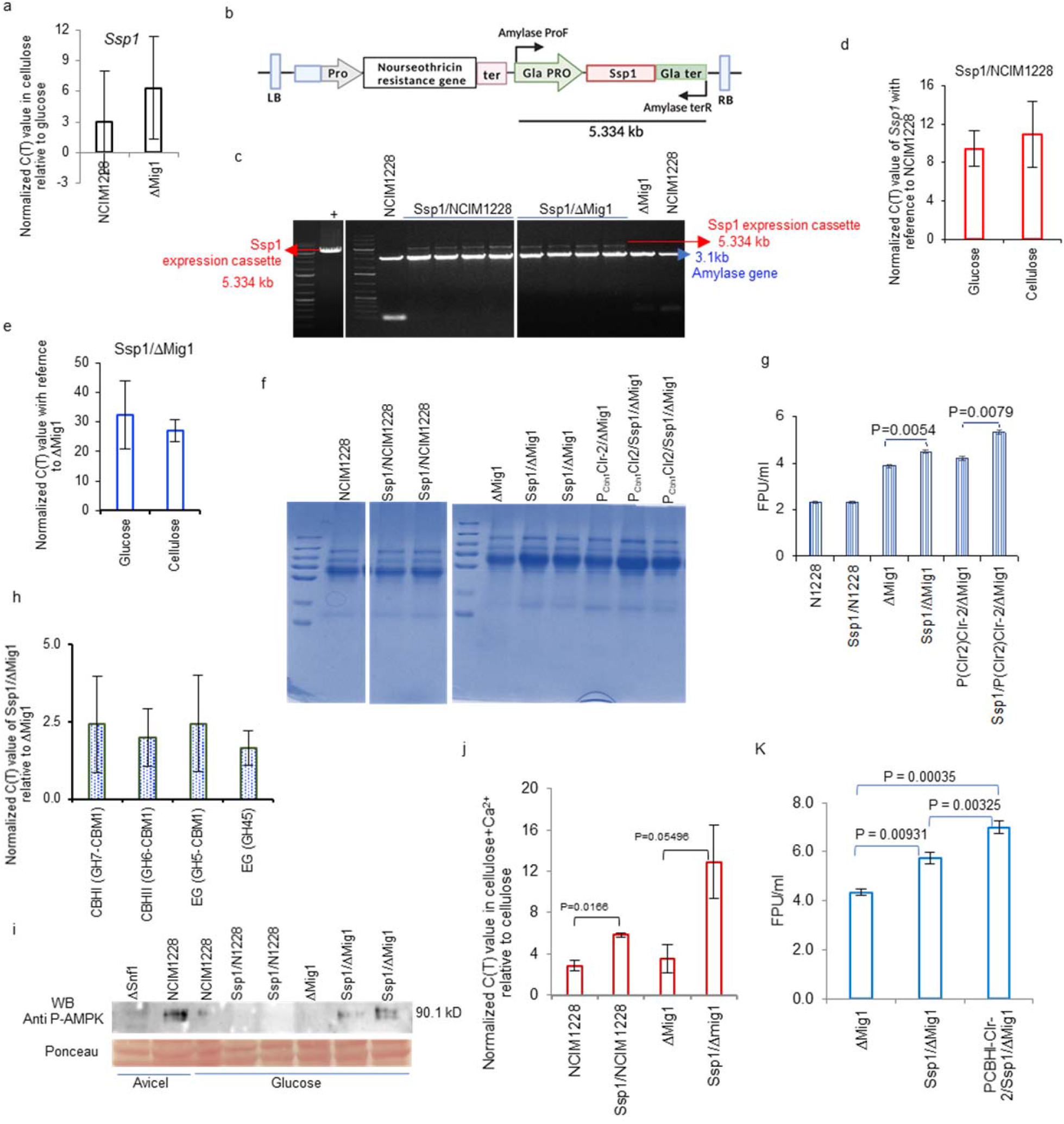
SSP1 activates SNF1 by phosphorylation. **(a)** An increase in transcript levels of SSP1 in NCIM1228 and ΔMig1 was determined by RT-PCR in cellulose relative to glucose. **(b)** Schematic of Sak1 expression cassette. **(c)** Confirmation PCR of Ssp1 expression cassette integration. Transcript levels of Ssp1 when Ssp1 was over-expressed in **(d)** NCIM1228 and **(e)** ΔMig1. **(f)** SDS-PAGE profile and **(g)** total cellulase activity present in the secretomes of parent and engineered strains grown in cellulosic RCM medium. **(h)** Transcript levels of major cellulases in Ssp1/ΔMig1 compared to ΔMig1 in MM medium having cellulose. **(i)** P-SNF1 AMPK Western blot showing P-SNF1 levels in NCIM1228, Ssp1/NCIM1228, ΔMig1, and Ssp1/ΔMig1 grown in glucose; whole cell extracts of ΔSnf1 and NCIM1228 grown in cellulose were taken as controls. **(j)** Transcript levels of *ssp1* in cellulose+Ca^2+^ relative to cellulose. **(k)** Total cellulase activity of Ssp1/ΔMig1, and P_Cbh1_Clr-2/Ssp1/ΔMig1 determined by FPU assay, when the strains were grown in RCM medium having 5g/l CaCO_3_.

We also examined the effect of Ca^2+^ on the *ssp1* transcript levels and cellulase production by SSP1/ΔMig1. We indeed found increased *ssp1* transcripts in cellulose+Ca^2+^ relative to cellulose (Fig. 6j); moreover cellulase activity determined by FPU assay was ∼6 FPU/ml in SSP1/ΔMig1 grown in cellulosic RCM medium having 5g/l CaCO_3_ (Fig. 7k). We also over-expressed *ssp1* in P_Cbh1_Clr-2/ΔMig1 and achieved ∼7FPU/ml in its secretome obtained from RCM medium having 5g/l CaCO_3_ (Fig. 7k). This confirms complementary role of SNF1 and Ca^2+^ signaling on cellulase production.

## Discussion

Our study elucidates that cellulose solely is sufficient to induce transcriptional activation of cellulase via Clr-2 in *P. funiculosum*, however twin signal of cellulose+Ca^2+^ is vital to cellulase translation and secretion (Fig. 6i). Further, increasing the transcript levels of *Clr-2* is effective only when the downstream Ca^2+^ signaling is also accelerated to achieve homeostasis between the cellulase transcription and translation. Moreover, we found extracellular secretion of cellulase to be a highly regulated and coordinated event during carbon stress. We identified the indispensable role of SNF1 AMPK as a Ca^2+^-activated kinase in cellulase secretion. We also identified CaMKK SSP1, which phosphorylates SNF1 in the presence of cellulose+Ca^2+^ during carbon stress. Since, SSP1 over-expression couldn’t increase P-SNF1 levels in NCIM1228, degradation of MIG1 during carbon stress seemed essential to maintain SNF1 in its phosphorylated form. The study suggests the crosstalk between MIG1 and SNF1 via SSP1 during carbon repression. Since role of SSP1 has previously been found in the cell proliferation and coordination of meiosis with spore formation in *S. pombe*, we believe that these processes are linked to cellulase production in filamentous fungi.

First, because exocytic secretion solves dual purpose of secreting cellulase to the exterior while propagating hyphal growth towards the crystalline cellulose, and second, cellulose being the most recalcitrant carbon source, must be the last available carbon source in the proximal environment and therefore might be a signal of the arrival of unfavorable conditions, inducing meiosis.

Quantitative proteomics of NCIM1228 and ΔSnf1, supported by functional studies aided in revealing the mechanistic of cellulase production by *Penicillium funiculosum*. Ca^2+^ signaling based upsurge of processes, like autophagy and proteasomal degradation, meet the simultaneous requirement of energy, cell proliferation, and cellulase production during carbon limiting conditions. Ca^2+^-signaling also activates conserved HOG1 MAPK pathway, to activate the expression of osmostress responsive genes. Osmolytes like glycerol and trehalose might be involved in protecting the cell from autophagic enzymes and oxidative stress caused by increased metabolic activity to produce cellulases. Activation of HOG1 blocks protein secretion and also inhibit transporters which might be helping the mycelia to retain osmolytes, as well as carbon and nitrogen sources from escaping^31,64–66^. Ca^2+^-activated SNF1 AMPK act as a balancing factor and determines an upper threshold for all calcium-activated processes including phosphorylation levels of HOG1 MAPK. Downregulation of P-HOG1 by SNF1 is essential as constitutively increased levels of P-HOG1 impede vesicular transport and fusion to the plasma membrane, the last steps of exocytic secretion, resulting in secretion block of cellulases.

SNF1 driven equilibration of the cellulase stimulating processes (proteasomal degradation, feeder pathways of central carbon metabolism, oxidative phosphorylation) and inhibiting processes (hyperphosphorylated Hog1) seems essential for the cellulase secretion, as even modest increase in hyperosmolarity imbalanced the delicate equilibrium of P-HOG1, impeding secretion. This also provides the reason of slow growth of ΔSnf1 on complex carbons. Studies on *Yarrowia lipolytica* have also shown the adverse effect of hyperosmolarity on protein secretion^68^. PTP2, a conserved phosphatase, regulates HOG1 MAPK phosphorylation during salt stress in fungi^69^. Recently, additional phosphatases (PTP1, PP2C, and CDC14) were also found to dephosphorylate HOG1 MAPK in response to stress^70^. It is yet to be seen if their activation is SNF1 dependent during carbon stress. We are therefore analyzing, if over-expression of these phosphatases could complement *Snf1* deletion in maintaining HOG1 in its dephosphorylated state during cellulase production.

Further, SNF1 is known to inhibit mTORC1 via Tsc complex, a conserved mechanism among eukaryotes^71^. Activated by RHEB (Ras-like small GTPase), mTORC1 favours anabolic processes upon glucose availability^71^. We identified elevated levels of RHEB in ΔSnf1 cellulose+Ca^2+^ proteome, in addition to high surge in mTORC1-regulated processes, e.g. ribosome and tRNA biogenesis, mRNA surveillance and amino acid biosynthesis. It indicated that SNF1 controls the mTORC1-regulated pathway in filamentous fungi via downregulating RHEB. Further analyzing the phospho-proteome in cellulose+Ca^2+^ vs cellulose in NCIM1228 and ΔSnf1 would reveal the complete regulon of cellulase production. Nevertheless, it is clear that SNF1 maintains homeostasis in the cell under carbon stress condition by controlling both HOG1- and mTORC1-mediated pathways, which subsequently support cellulase enzyme secretion outside the cell.

To conclude, our study highlights segregated control of cellulase transcription, post-transcription and secretion by Clr-2, Ca^2+^-mediated signaling, and Snf1 AMPK, respectively. Supported by various studies in diverse yeasts and fungi, the mechanism seems conserved for the secretion of enzymes involved in the hydrolysis of other nonfermentable carbon sources.

## Methods

### Strains, plasmids, growth media and culture conditions

All strains, plasmids and primers used in this study are listed in Supplementary Table 1, 2, and 3 respectively. All fungal strains are derivatives of *P. funiculosum* NCIM1228. Gene deletion and over-expression in *P. funiculosum* were carried out as previously described^35^.

The growth medium used in the study are Potato dextrose (Himedia) broth for conidiospore germination, yeast nitrogen base with ammonium sulfate (Difco) agar for growth profiles, low malt extract peptone (LMP) agar^35^ for transformations, Mandel’s medium (MM) for transcriptomic, RNA-profiling by RT-qPCR and proteomics studies, and RCM/MM medium for cellulase production^11,12^.

Primary cultures are prepared by adding 10^6^ conidiospores to the 50 ml PD broth and cultures are allowed to grow at 28ºC for 30 hours at 120rpm. Secondary cultures are inoculated with 10% primary cultures. Growth profile on agar plates is observed by spotting 5μl of 10^4^ conidiospores/ml suspension on SC agar with 2% carbon source followed by 48 hours of incubation at 28ºC. For determining differential growth profile in liquid medium, Mandel’s medium with 4% glucose is inoculated with primary culture at 5% and the cultures are incubated at 28ºC for 24 hours. Mycelia is collected by filtration through Mira cloth and the collected mycelia is dried for 72 hours at 70ºC and weighed.

For cellulase induction and production, RCM having 2.5% CaCO_3_ or Mandel’s medium having CaCl_2_ (50mg/l) CaCO_3_ is used unless specified otherwise and the experiments describing no calcium have no known source of Ca^2+^ added to RCM or Mandel’s medium. For transcriptomics and proteomics studies, strains are grown in Mandels medium for 48 hours and mycelia was collected at logarithmic phase and processed accordingly.

### Enzyme Assays and Biomass hydrolysis

Secretomes are evaluated for enzyme activities on 0.5% crystalline cellulose (Avicel, Sigma), 1% carboxymethyl cellulose (CMC, Sigma), 1% xylan (Himedia), and 1% starch (Himedia) as described previously^11^. Citrate–phosphate buffer (50 mM, pH 4.0) is used for diluting the secretomes and carrying out enzyme assays. Enzyme and substrates are incubated together for 30 min (2h for crystalline cellulose) at 50°C. The sugars released by enzyme action are measured by dinitrosalicylic acid (DNSA) method. The absorbance at 540 nm is measured relative to glucose or xylose standard curve. One unit of enzyme activity is defined as the amount of enzyme releasing 1 μmol of reducing sugar per min. *p*-nitrophenyl-β-d-glucopyranoside (pNPG) (sigma) is used as substrate to determine β-glucosidase activity in the secretome and the amount releasing 1 μmol of *p*-nitrophenol per min is considered one pNPGase unit. The absorbance of test at 410 nm is measured relative to *p*-nitrophenol standard curve. Additionally, β-glucosidase activity is also detected by zymogram technique using 4-Methylumbelliferyl-beta-D-glucuronide hydrate (4-MUG) (sigma) as substrate. Secretomes are electrophoresed on 10% Native PAGE; electrophoresed gels are rinsed with water and Citrate–phosphate buffer (50 mM, pH 4.0), followed by incubation in 5 µM 4-MUG solution for one hour. The zymogram gels are visualized under UV light to detect fluorescence due to the 4-methylumbelliferone (4-MU) released. followed by Total cellulase activity of secretome is measured by filter paper assay that measures fixed degree of conversion of substrate as previously described^11^. One filter paper unit is the amount of secretome that releases 2 mg glucose from 50 mg filter paper in 60 min at 50°C.

Based on the protocol discussed in Olusola et al (2020a), secretome were evaluated on nitric acid pretreated rice straw using two substrate loading concentrations (15 % and 20 % dry biomass) at enzyme concentrations of 30 g/kg DBW (3% enzyme load). Hydrolysis reaction was conducted in 1.2-ml 96-well plates having pretreated biomass at a 15 % and 20 % dry weight loading in a 250μl final reaction volume. The protein content of the secretomes were measured, and the appropriate volume having 30 g/kg DBW was added to the reaction mixture. The reaction was conducted in 50 mM citrate-phosphate buffer (pH 4.0) and incubated at 50 °C with constant shaking at 200 rpm for 72 h. Control experiments were carried out under the same conditions using substrates without enzymes (enzyme blank) and enzymes without substrates (substrate blank); a substrate-free negative control was set up by filling wells with 50 mM citrate-phosphate buffer, pH 4.0, and the background of soluble sugars present in the respective biomass was determined by incubating each biomass in the absence of enzymes. Following the completion of hydrolysis, the plates were centrifuged at 3000g for 10 min in a swinging bucket centrifuge (Eppendorf, Germany) to separate the solid residue from the digested biomass. Hydrolysates were analysed for their sugar content by high-performance liquid chromatography equipped with an Aminex HPX-87H anion exchange column (Bio-Rad, USA) and a refractive index detector and the percentage release was calculated citing the theoretical yield.

### Confocal microscopy

*P. funiculosum NCIM1228* was grown in Mandel’ medium with 4% cellulose with and without calcium for 48 hours. Mycelia was collected by centrifugation at 3000 rpm for 5 minutes and washed with PBS three times. Mycelia was incubated in solution containing 1µM ER-tracker, 4µM DAPI and 2µg/ml Bodipy at RT for one hour. Stained mycelia was washed with PBS three times and suspended in PBS before being observed under a Nikon A1R confocal microscope. One way Anova test was performed for morphological differences with in the sample and t-tests were performed to test the significance between two test conditions.

### Transcriptomic studies by RNA-seq

NCIM1228 and ΔMig1 are grown in Mandel’s medium having 4% glucose/Avicel. Total RNA is isolated from log phase cultures by Qiagen Plant mini RNeasy kit according to the manufacturer’s instructions. Traces of DNA, if any, are removed by DNase treatment before proceeding for RNA sequencing. RNA-seq is carried out on HiSeq 2000 platform with 125×2 paired-end read chemistry (Bionivid Technology Pvt Ltd). Biological replicate sequencing libraries for both strains are created with poly-A tailed mRNA enrichment using the standard Illumina TruSeq mRNA RNA-Seq protocol. RNA-Seq reads obtained are assembled using Trinity with reference genome guided approach. Assembled transcripts are quantified by mapping generated sequencing reads to the assembled transcripts using the alignment mapping program Bowtie2 and alignments are coordinate-sorted by SAMtools. Quantitative program RSEM generates fragments per kilobase of transcript per million mapped reads (FPKM) from the quantitated data. The protein domains are predicted using InterProScan. For differential expression profiling, all FPKM values are normalized to the library size using the R package, Edge R. The obtained p-values are plotted against log_2_ fold change using Volcanose R and volcano plots are obtained to assess the significance of up- and downregulation of transcripts as shown in respective figures. The DNA binding and Calcium activated proteins are filtered and identified from InterPro result. Heatmaps are generated from log_2_ FPKM values of selected transcription factors and calcium activated proteins by performing Hierarchical Clustering using the Euclidean distance matrix option of morpheus heatmap tool.

### Expression profiling by real-time qPCR

For real-time PCR experiments, mycelia is harvested from log phase cultures and total RNA is extracted using RNeasy kit (Qiagen). Traces of DNA are removed by DNase (Invitrogen) treatment prior to cDNA synthesis and RNA concentration is measured by nano-drop. Equal amount of RNA is used to synthesize cDNA by Invitrogen cDNA synthesis kit. cDNA is used as template to carry out qRT-PCR using iTaqTM Universal SYBR^®^ Green Supermix (Bio-Rad) and Bio-Rad CFX96 qPCR detection system. qRT-PCR is done in biological triplicates with tubulin as the endogenous control. Relative expression levels are normalized to tubulin, and fold changes in RNA level are the ratios of the relative expression level of PfMig188 to NCIM1228 under repressing conditions and cellulase inducing conditions to no carbon conditions.

### Construction of gene over-expression and deletion cassettes

#### *Ctf1b expression* cassette

A 3953bp DNA fragment having *Ctf1b* gene along with 1000bp promoter and 200bp terminator was PCR amplified from genomic DNA using primers Ctf1b Pro EcoRI F and Ctf1b Ter MluI R and cloned in pBIF replacing T-DNA at *Eco*RI and *Mlu*I restriction sites. As a result, 12.976 kb binary vector was created having *Ctf1b* expression between the T-DNA arms.

#### Clr-2 over-expression cassettes

Clr-2 cassette (1005bp promoter+2753bp Clr-2gene+207bp terminator) was amplified from NCIM1228 gDNA with Clr2PF1 and Clr2R2 primers. The fragment was cloned in pBIF at *Mun*I/*Bam*HI restriction sites, replacing Gpd promoter and GFP gene with Clr-2 cassette in T-DNA of pBIF. For Clr-2 under Gpd promoter, 2753bp Clr-2gene+207bp terminator was amplified from NCIM1228 gDNA with Clr2F1 and Clr2R2 primers, and cloned at *Sac*I/*Bam*HI restriction sites. For Clr-2 expression under CBHI promoter, CBHI promoter was amplified from NCIM1228 genome using Cbh1Pro-F and Cbh1 Pro-R and cloned in pBIF at *Eco*RI*/ Sac*I restriction sites replacing Gpd promoter, followed by Clr-2 cloning at *Sac*I/ *Bam*HI restriction sites.

#### *Snf1* disruption cassette

A DNA fragment of 2620 bp having Snf1 gene with flanking regions of 500bp was PCR amplified from genomic DNA using primers Snf1 XhoI F and Snf1 MauBI R and cloned in pCambia1302 at *Xho*I and *Mau*BI restriction sites, resulting in pSnf1. A 10014 bp binary vector having Snf1 disruption construct, was generated by removing 639 bp Snf1 ORF region from pSnf1 by restriction digestion and replacing it with 1414-bp zeocin resistance cassette at *Spe*I/*Hind*III restriction sites.

#### Sak1 over-expression cassette

First, 1026 bp hygromycin resistance gene (*hph*) was replaced with 573 bp nourseothricin resistance gene (*Nat*) at *Aat*II/*Eco*RI restriction sites in pCambia1302 resulting in pNat. Ssp1 gene flanked by glucoamylase promoter and terminator was chemically synthesized and the resultant cassette of 5317bp was cloned into pNat at *Eco*RI/*Mau*BI restriction sites.

### Western blot analysis

CBH1 levels in the secretome was detected by western blotting as previously described^34^. For detection of intracellular CBH1 mycelial extracts are collected by filtration using mira cloth, washed three times with excess of distilled water, and extra moisture was removed by pressing between the layers of filter paper before lyophilization. 100 mg of finely grounded lyophilized mycelia is added to 1ml of lysis buffer (50mM Tris-Cl, pH8.0, 0.05% SDS, 0.1% sodium deoxycholate, 0.1% Triton X-100, 5mM Sodium pyrophosphate, 50mM Sodium fluoride, 0.1 mM sodium vanadate, 0.05% PMSF, cOmplete™ Mini, EDTA-free Protease Inhibitor Cocktail (Roche), and phosSTOP (Roche)) in bashing bead lysis tubes (Zymo Research) and cells are lysed by 30 lysis cycles (1 min bead beater and 1 min ice). Mycelial extract is collected by centrifugation at 13000 rpm fo r30 minutes. The whole mycelial extract is evaluated for its protein content by BCA method. 50µg of whole mycelial protein is electrophoresed on 12% SDS-PAGE, the electrophoresed proteins are transferred to nitrocellulose membrane using semi-dry blot technique. For western blotting, the membranes are blocked with 5% BSA in Tris buffer saline with 0.1% tween-20 (TBS-T) for 2 hours, prior to incubation with rabbit Anti-CBH1(1:5000) overnight with regular shaking. The blots are washed with TBS-T three times and incubated with HRP conjugated anti-rabbit/mouse for one hour. The blots are developed by G-biosciences ECL reagent and chemiluminescence is detected using high-resolution chemiluminescence mode on Bio-Rad chemidoc XRS+ system. Similar procedure is followed for western blotting with Anti-phosphoSnf1 AMPK and Anti-phospho Hog1 MAPK.

### Quantitative proteomics by LC-MS/MS

5 ml of primary culture of NCIM1228 and ΔSnf1 were added to Mandel’s medium having cellulose (4%) and cellulose (4%) **+** calcium (0.5%) in triplicates. MM cultures were incubated at 28°C with regular shaking at 120rpm. A part of the culture was used to collect mycelia at 48 hours of growth, however the remaining culture fraction was incubated for another 72 hours to harvest the secretome. Before proceeding with the mycelial proteomics sample preparation, the secretomes were examined for the desired phenotype by SDS-PAGE and enzyme assays. After confirming the distinct phenotype of the two strains under given conditions, whole cell proteome was extracted from mycelial samples and quantified by BCA method. Next, we added 1µg ^13^C and ^15^N labelled human Apolipoprotein (Apo-1) (Sigma) as a known standard to 50µg total mycelial extract to assess the variance during sample preparation and measurements. Whole mycelial extracts with Apo-1 standards were alkylated, and reduced prior to proteolytic digestion with trypsin (1µg/50µg of protein) for 16 hours at 37°C. Peptides were purified and concentrated by C18 spin columns (Thermo Fisher Scientific). Purified samples were vacuum-dried and reconstituted in 0.1% (v/v) formic acid before before proceeding for absolute quantification of all proteins by LC-MS/MS analysis by Thermofisher Orbitrap Fusion™ Lumos™ Tribrid™ Mass Spectrometer equipped with nano-LC Easy nLC 1200.

Liquid chromatography separation was performed at a flow rate of 300 nL/mL on a C18 pre-column (Acclaim PepMapTM 100, 75 μm×2 cm, nanoViper, P/N 164946, ThermoFisher Scientific Incorporation) followed by analytical column (Acclaim PepMapTM RSLC C18, 75 μm×50 cm, 2 μm, 100 Å, P/N ES803). The peptides were separated using a gradient of solvent B (95% acetonitrile in 0.1% formic acid) from 2-10% in 5 min, gradient increase to 45%, followed by sharp increase to 95% for 10 min. The sample is injected into the mass spectrometer and the MS1 data is acquired by Thermo Xcalibur software setup version 4.3.73.11 (Thermo Fischer Scientific, Inc 2019) in full scan mode at 120,000 orbitrap resolution with mass range from 375 to 2000 Da. The precursors are fragmented using Higher-energy C-trap dissociation (HCD) in ion trap (IT) detector with collision energy of 28 in a data dependent MSN Scan acquisition. Charge state screening of precursor ions and monoisotopic precursor selection was enabled. The parent ions once fragmented were excluded for 40 s with exclusion mass width of ± 10 ppm. The lock mass option (polydimethylcyclosiloxane; m/z 445.120025) enabled accurate mass measurement in the MS Mode.

For analysis, raw LC–MS/MS data files obtained from the mass spectrometer were processed with Proteome Discoverer™ (Version 2.4.1.15, Thermo FisherTM Scientific Inc.). For the search process, Mascot and Sequest HT tools were used. Peak lists obtained from MS/MS spectra were identified using the MSF files. Protein identification was conducted against a concatenated target/decoy version of the in-house predicted proteins or in-house database (11,213 target sequences) obtained from the draft genome sequence of *Penicillium funiculosum*, and Apo-1 protein sequence (Sigma) along with Proteome Discoverer™ contaminant database. The identification settings were as follows: trypsin digestion with maximum of 2 missed cleavages; minimum peptide length 6; precursor mass tolerance 20 ppm; fragment mass tolerance 0.5 Da; fixed modifications; carbamidomethyl c (+57.021464 Da), variable modifications; oxidation of m (+15.994915 Da), acetylation of protein n-term (+42.010565 Da). Peptides and proteins inferred from the spectrum results using SwissProt and Uniprot database. Peptide Spectrum Matches (PSMs), peptides and proteins were validated at a target False Discovery Rate (FDR) strict to 0.01 and relaxed to 0.05. Mycelial protein abundance data was normalized to the labelled peptide abundance data before quantification. The formula for normalizing the sample is as follows:

x = x_1_xx_2_

x = abundance of a given protein

x_1_= true abundance of a protein in a given sample;

x_2_= variability due to sample processing and data acquisition

In case of Apo-1,

x_Apo-1_ = x_2,_ as x_1_ is constant i.e. (amount of Apo-1 added to each sample is 1µg).

Sum total Abundance of Apo-1 in 8 samples = Abundance of 8 µg Apo-1,it can be written as 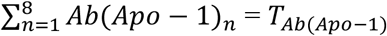 where *T*_*Ab*(*Apo*-1)_ represents total abundance of 8µg of Apo-1

Now, x2 = *Ab*(*Apo* − 1)_*n*_ × *ν*_*n*_, ∀*n* ∈ [1,8]; where v_*n*_ is variability factor of sample n

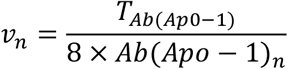

and we achieved

*Ab*(*Apo* − 1)_1_ × *ν*1 = *Ab*(*Apo* − 1)_2_ × *ν*2 = *Ab*(*Apo* − 1)_3_ × *ν*3 = *Ab*(*Apo* − 1)_*n*_ × *νn* We assumed that all the proteins in a given sample will acquire variation during sample processing and data acquisition. Variability factor of Apo-1 would be applicable to all proteins processed together in a given sample, therefore multiplying the protein abundance of all proteins with variability factor of Apo-1 in a given sample will nullify the effect of variation due to sample processing. Normalized protein abundance values of NCIM1228 and ΔSnf1 secretomes containing cellulose was compared to the secretome with cellulose + calcium by ANOVA (p<0.05) and student’s t test (p<0.05). To evaluate the protein distribution pattern under each of the test conditions, data was subjected to change in the abundance level of a particular protein across all samples Anova test.

A total of 129231 peptides were detected at a 1% FDR on the peptide spectrum match, representing 5567 proteins among 12 samples. A two-way ANOVA of normalized protein levels across four groups was performed, and samples were verified for variance among different groups and similarity between the replicates. Out of the three technical replicates, one of the technical replicates of ΔSnf1 grown in Avicel + Ca^2+^ showed drastic variation from the other two, and was therefore ignored. For the same reason, we used two technical replicates for each condition for further data analysis. We set the threshold at 2 unique peptides per protein and, also ignored all the proteins with gaps which were not represented in the whole sample group. A total of 2979 proteins represented in all the samples by at least 2 unique peptide were used for further analysis. Principal component analysis was performed using XL STAT trial version.

### Antifungal susceptibility assay

200 µl of *P. funiculosum* NCIM1228 primary culture was added to 10 ml Mandel’s medium having different concentrations of SHAM, boscalid, epoxiconazole, fluazinam and metalaxyl in triplicates for each concentration. The cultures were incubated at 28°C for 24 hours at 120 rpm and mycelia was collected by filtration through mira cloth. Mycelia was dried at 70°C for three days and weighed. EC50 of all antifungals was determined for *P. funiculosum* NCIM1228 by dry mycelial weight at different concentrations compared to control. Next, 200 µl of primary cultures of NCIM1228 and ΔSnf1 were added to 10 ml of Mandel’s medium having Antifungals at their EC_50_ concentrations in triplicates along with MM having no antifungal as control. Difference in growth patterns of NCIM1228 and ΔSnf1 was determined by percentage growth at EC_50_ compared to control cultures.

### Statistical analysis and online tools and databases

We used Microsoft excel, and Graphpad-prism for creating graphs, for performing unpaired t-tests with Welch correction and one way and two way ANOVA. PCA was performed using XL-STAT trial version. Online web applications, VolcanoseR^72^, Morpheus (https://software.broadinstitute.org/morpheus), and Interactivenn^73^ were used to create volcano plots, heatmaps, and venn diagram respectively. Figures were created using Microsoft PowerPoint and bioRENDER (Fig.6). We used BLAST for homology based search, SMART for identifying domain boundaries of Snf1 AMPK, and STRING database with *Penicillium marneffei* as the query organism to identify interactome of putative proteins of *P. funiculosum* NCIM1228. All NCIM1228 proteins used as query showed more than 80% similarity to respective *P. marneffei* proteins.

## Data availability

Any additional information required to reanalyze the data reported in this study is available from the lead contact upon request.

## Authors’ contribution

AR and SSY designed experiments; AR conducted experiments on transcriptomics and proteomics, and performed computational analysis; AR, OAO and KJ did cloning; AR and OAO performed fungal transformations; AR and MD carried out enzyme assays, AR performed western blotting of intracellular P-SNF1 and P-HOG1, AR and TS performed western blotting of secreted and intracellular CBH1. AR performed chemical genetic experiments; AR, and SSY wrote manuscript. All authors read and approved the final manuscript.

## Acknowledgements

We thank Prof Alok K Mondal, Jawaharlal Nehru University, New Delhi, for providing Anti - Phospho-p38 antibody used for the western blotting experiments. We acknowledge Girish H Rajacharya for managing mass spectrometry facility at ICGEB, New Delhi. We acknowledge Dr. Kanu Goel, AMITY University, Punjab for her guidance in arriving at the mathematical equations used in the proteomics data analysis. AR was supported by DST-SERB, India via national postdoctoral fellowship (PDF/2018/002549). This work was funded by Department of Biotechnology, Government of India via Bioenergy Centre grant no. BT/PR/Centre/03/2011-Phase-II.

